# Impacts of sea level rise on an endemic butterfly and its freshwater wetland habitat in the Florida Keys

**DOI:** 10.1101/2025.06.16.659612

**Authors:** Sarah R. Steele Cabrera, Robin Sarabia, Erica H. Henry, Chad T. Anderson, Jaret C. Daniels

## Abstract

Coastal and island ecosystems are disproportionately vulnerable to sea level rise and other impacts of anthropogenic climate change. Using monitoring data collected between 2013 and 2024, we explore habitat shifts, population dynamics, and phenology of an endemic butterfly in the Florida Keys, Klot’s sawgrass skipper (Euphyes pilatka klotsi). Range-wide surveys of the butterfly and its associated habitat demonstrated that the butterfly’s sole host plant is decreasing in abundance at lower elevations, likely due to increased salinization. We also find declines in butterfly population size over an eleven-year period using distance sampling. Despite these declines, we find that Klot’s sawgrass skipper’s range has not contracted significantly over the study period.

This study demonstrates that monitoring a single taxon and its host is useful for understanding how sea level rise impacts a keystone plant species in freshwater wetlands which are vital for the continued survival of a suite of rare and endemic species found in the Florida Keys. Projections of near-future sea level rise indicate that most or all of this habitat will be lost within several decades; the continued study of low-lying islands is critical to gain insight into the global phenomenon of sea level rise.

## Introduction

Sea level rise (SLR) is one of the most certain outcomes of anthropogenic climate change, disproportionately impacting coastal ecosystems [1–3]. Current global projections of SLR by 2100 range from 0.28-1.01m [4]. Low-lying islands are particularly vulnerable to SLR [5] and also have disproportionately high levels of biodiversity and endemism [6]. Given the importance of SLR in the coming decades, it is imperative to understand how this phenomenon is impacting ecosystems and biodiversity.

Insects are the most diverse group of animals [7] and are vital in both natural and human-dominated systems for functions including nutrient cycling, pollination, and as food for other taxa [8,9]. Despite this, they are vastly understudied, particularly in tropical regions where biodiversity is highest [7,10], hampering efforts to understand how insects are being impacted by global climate change and other contemporary factors threatening biodiversity [11]. Predicting winners and losers under global climate change is challenging and often trait-[12,13] and/or species-specific [14]. In general, populations of species with narrow ranges, on mountains [15] and islands [16] are more likely to decline or go extinct, but these trends are not universal [17].

Butterflies are often used as indicator species because of their reliance on specific resources during their multi-phasic life cycles [18,19]. Like other insects, they are highly sensitive to react quickly to environmental change [20]. Although the predictive value of bioindicators is variable [21] and not useful in all situations [22], we demonstrate that monitoring one butterfly taxon and its host plant provide informative indicators of ongoing changes occurring in an ecological community on the front lines of impact by climate change. We explore habitat shifts, population dynamics, and phenology of an endemic butterfly in the Florida Keys, Klot’s sawgrass skipper (Euphyes pilatka klotsi) to understand how freshwater wetlands are changing in response to increased salinization due to climate change.

## Methods

### 2.1 Study Organism

Euphyes pilatka klotsi L. Miller, Harvey and J. Miller, 1985, also known as Palatka skipper, Klot’s skipper, or sawgrass skipper, is a large rusty-brown grass-skipper [23] (Figure 1) endemic to the Lower Florida Keys [24] (Figure 2); E. p. pilatka is found across much of the coastal southeastern US from Virginia to Mississippi [23]. Klot’s sawgrass skipper has been observed on nine islands in the Lower Florida Keys (From East to West: No Name, Big Pine, Little Torch, Middle Torch, Big Torch, Ramrod, Summerland, Cudjoe, and Upper Sugarloaf Keys, Stock Island) though it has apparently been extirpated from both the easternmost and westernmost extremes of its range [25,26]. It is currently extant from Big Pine Key to Upper Sugarloaf Key. Like its more widespread sister subspecies, Klot’s sawgrass skipper has a single larval food plant, swamp sawgrass (Cladium jamaicense Crantz), a sedge which is found in freshwater marshes and limestone depressions [27]. Eggs (Figure 1A) are laid singly on the host sedge and larvae create shelters comprising multiple sedge blades adhered with silk [28] (Figure 1B and 1C). Larvae have been observed to create multiple shelters as they grow, pupating inside of the final shelter.

**Figure 1:**
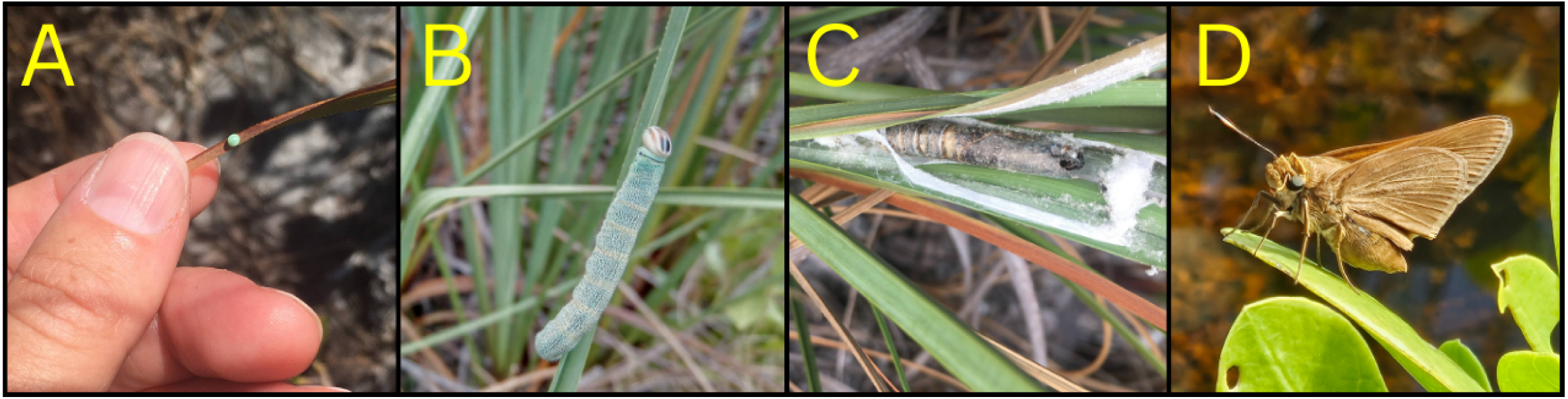
Life stages of Klot’s sawgrass skipper (Euphyes pilatka klotsi). A. Eggs are light green and laid singly on blades of their host plant, sawgrass. B. Larva on a blade of sawgrass outside of its shelter. C. Shed pupal tissue inside of a leaf shelter opened after the butterfly emerged. D. Adult perching on buttonwood (Conocarpus erectus)

**Figure 2:**
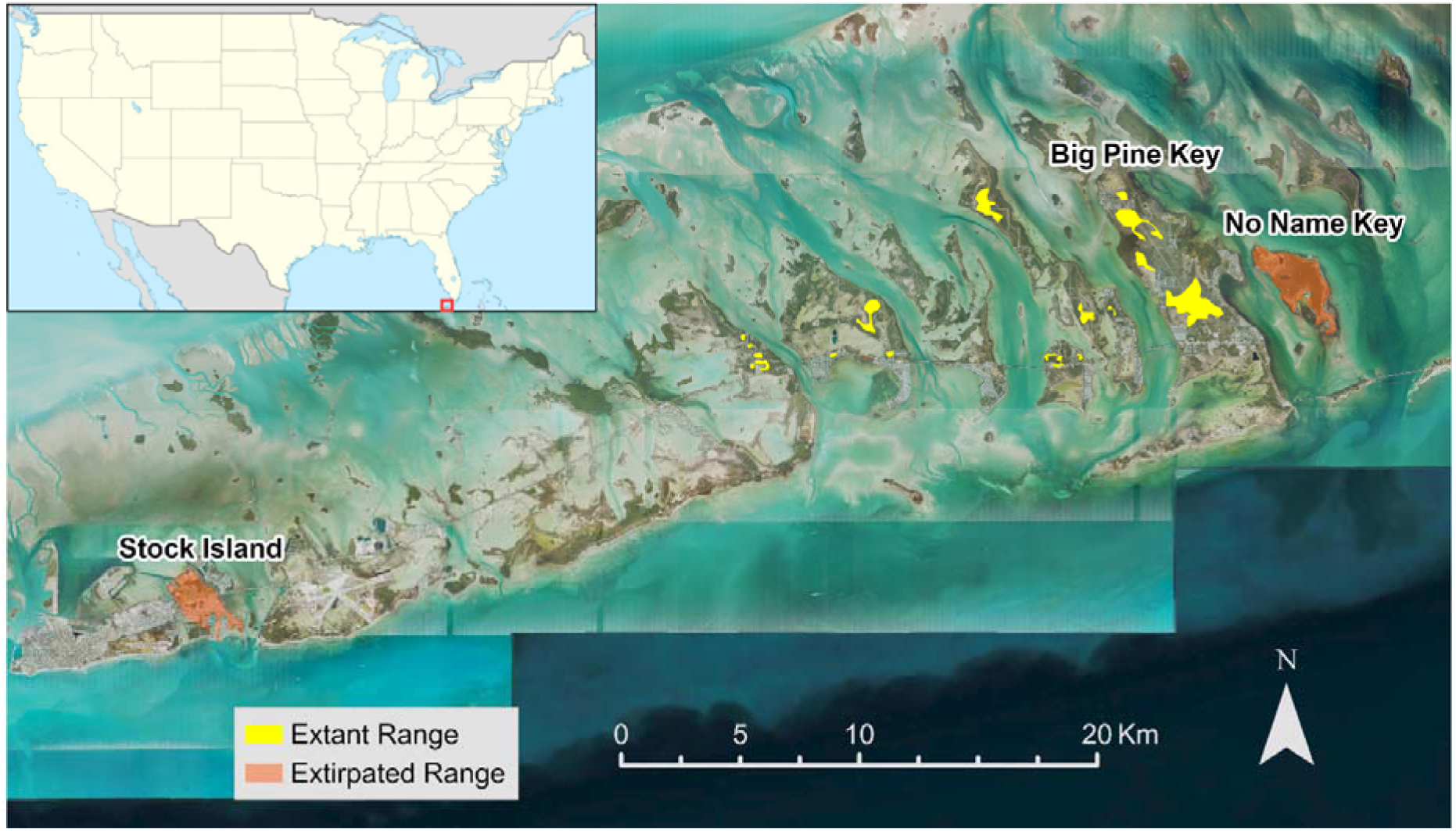
Map of the study area including the extant range of Klot’s sawgrass skipper and islands where this taxon has been apparently extirpated.

### 2.2 Study Site

The Florida Keys are a 250-km long archipelago located on the southern tip of Florida, USA which are composed of porous limestone derived from Pleistocene coral reefs [29]. Our study focuses on several islands in the lower Florida Keys (Figure 2), where the maximum natural elevation is approximately 2 m above sea level [30]. Several of these islands contain freshwater lenses that contract and expand with seasonal recharge, with most of the 100 cm of annual rainfall occurring between May to October [31]. These seasonal precipitation patterns are the only sources of freshwater input for island aquifers [32] and fresh water on these islands is subject to salinization from hurricanes and sea level rise [33]. Fresh water volume is further reduced by an estimated 20% in many areas due to a man-made canals [34]. Much of the terrestrial flora and fauna in the lower Florida Keys is directly dependent upon these limited freshwater resources, including a suite of endemic taxa that include well-known mammals like the Key deer (Odocoileus virginianus clavium), but also a number of plants and invertebrates [35,36].

### 2.3 Field Methods

We first located and mapped all potential Klot’s sawgrass skipper habitat. The entire land area categorized as freshwater habitat on public lands was extracted using 2009 Florida land cover data [37]. Additional small wetlands not included in the county land cover dataset were delineated manually from aerial photographs. A grid of 50-meter-long transects was created throughout all potential habitat from Big Pine and Sugarloaf Keys using ArcGIS 10.0 (Esri Redlands, CA, USA); in 2021-24 survey transects were 100 meters long. The selected habitat was then ground-truthed by trained observers who discarded any transects from which the host plant species was absent or that were unsafe or inaccessible to human surveyors. In 2013-2022, transects surveyed were chosen randomly. In 2024, surveys were limited to Big Pine Key due to limited personnel availability.

Field surveyors worked in a two-person team in March 2013 and independently in subsequent time periods. Between 2013 and 2016, surveys were conducted in Spring (March and April) and Fall (October) to coincide with presumed flight periods for adult butterflies based on the phenology of the mainland subspecies [38]. Surveys were conducted continuously between February 2021 and February 2022. A small number of surveys were conducted in November 2024. Factors such as a lack of available personnel and an extended government shutdown in October 2013 prevented additional data collection.

Surveys were conducted during daylight hours, between two hours after sunrise and two hours before sunset on fair weather days. Surveys were suspended during rain and high winds (above 15 knots). Trained surveyors traversed randomly selected transects on foot, using a digital map to minimize deviation from the transect. Upon encountering Klot’s sawgrass skipper, the observer recorded the organism’s distance from the transect when it was first observed to the nearest half meter, using a measuring stick to minimize error. Vegetation data was collected along butterfly survey transects in 2013 and 2021. Percent cover of sawgrass and shrub was estimated using Daubenmire cover classes [39] in 2.5m-radius circular plots along each transect at the beginning, middle, and end points of transects.

### 2.4 Statistical Methods

All analyses were conducted in R Version 4.4.1 [40]. To examine whether sawgrass and shrub abundance have changed over time, we subset the vegetation data to include only paired data where the same point was surveyed both in 2013 and 2021. We conducted two Wilcoxon signed-rank tests with continuity correction to approximate a continuous distribution on both the sawgrass and shrub data. To understand the effect of elevation on sawgrass abundance over time, we conducted two generalized least square regression models, one for the 2013 and one for the 2021 data, accounting for localized and exponential spatial autocorrelation using the nlme package [41] and LiDAR-derived elevation data [42].

To model butterfly detection and subsequently butterfly density estimates, we used the Distance package [43]. To improve model fit, butterfly detections were grouped into bins by distance. Each flight period was analyzed independently using the hazard-rate model with Hermite polynomial adjustment (selected by AIC), pooling observations across all islands. See Supplementary Table 1 for a comparison of all models and detection functions for each flight season. The resulting density estimates and standard errors were then compared to one another using pairwise t-tests.

## Results

We compared vegetation at a total of 210 paired points. We found that sawgrass abundance did not change over time (V = 2961.5, p = 0.3518). However, abundance values clustered more tightly around the mean in the 2021 data than in the more variable 2013 data, with fewer sampling plots with high sawgrass abundance (Figure 3, left panel). Shrub abundance decreased (V = 3567.5, p = 0.01214) between 2013 and 2021 (Figure 3, right panel).

**Figure 3:**
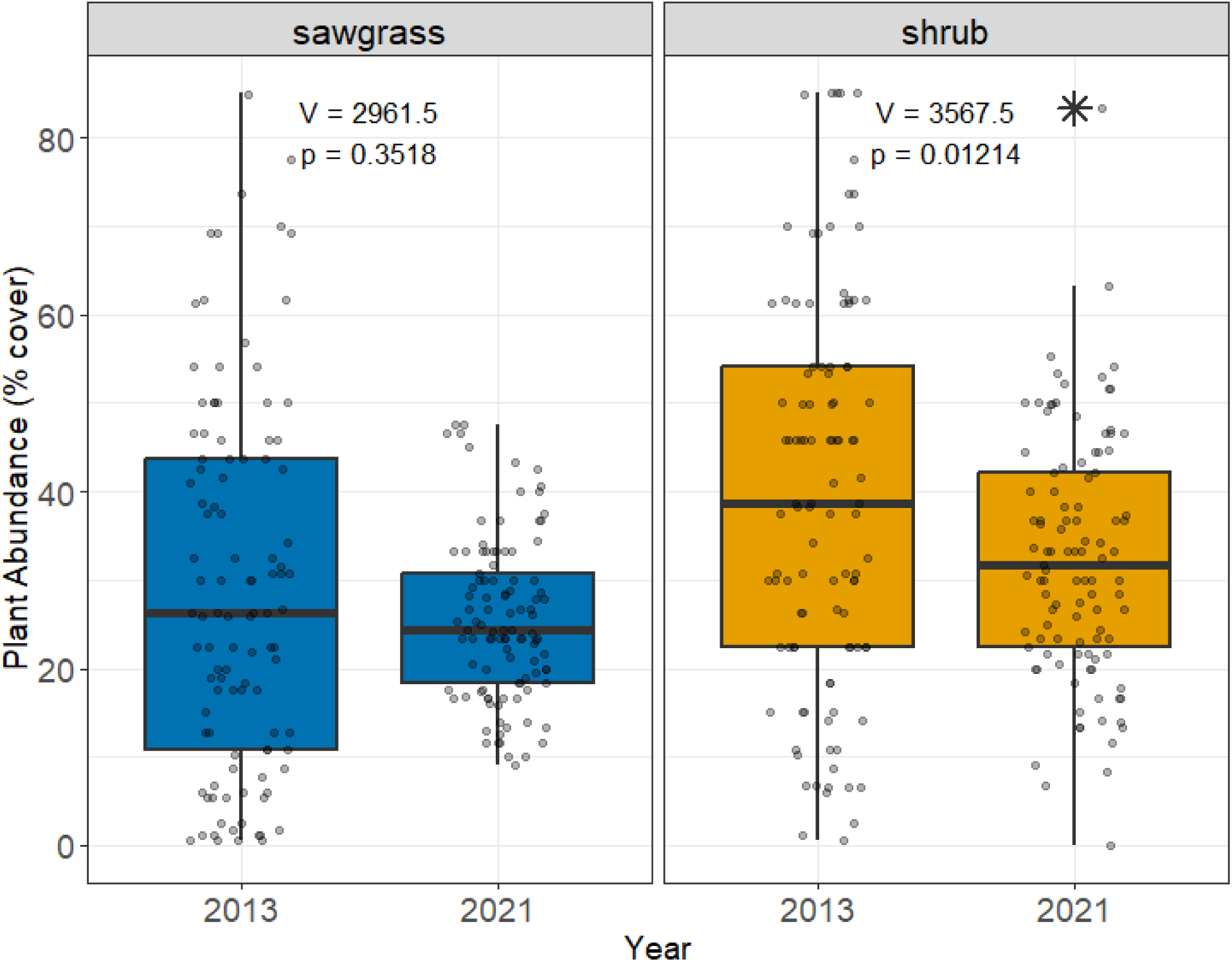
Boxplots comparing sawgrass (left panel) and shrub (right panel) abundance in 2013 and 2021 with test statistics for Wilcoxon signed rank test with continuity correction. Outlier data point is represented by an asterisk.

In the 2013 data, we found no relationship between sawgrass abundance and elevation (t = 0.247, p = 0.797; Figure 4 left panel). In 2021, we found that sawgrass abundance was positively associated with elevation (t = 3.208, p = 0.0014; Figure 4 right panel).

**Figure 4:**
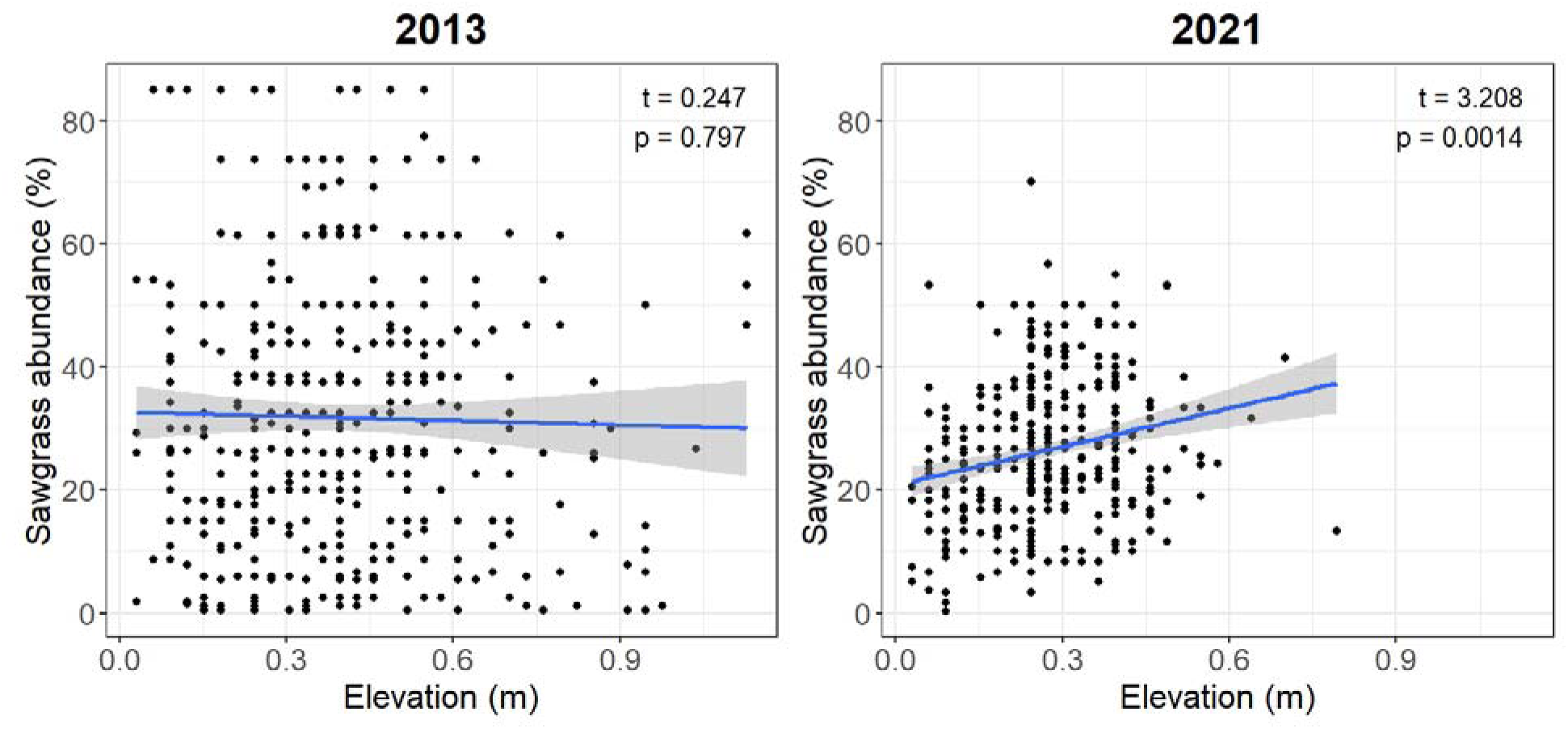
Scatter plots of sawgrass abundance by elevation in 2013 (left) and 2021 (right). Line represents the generalized linear model with the 95% confidence interval. Annotated statistics are from the generalized least square model which accounts for spatial autocorrelation in the dataset.

Butterfly population estimates in 2013-14 ranged from between 18.8 and 52.8 butterflies per hectare. In 2021-24, population estimates ranged from 0.4 to 4.6 butterflies per hectare (Figure 5). Estimates for two flight seasons (Fall 2014 and Spring 2021) had distributions that significantly deviated from the expected hazard-rate distribution (Supplementary Table 2). Estimates could not be generated for data collected in 2015 or 2016 due to limited sampling and the resulting small number of observations.

**Figure 5:**
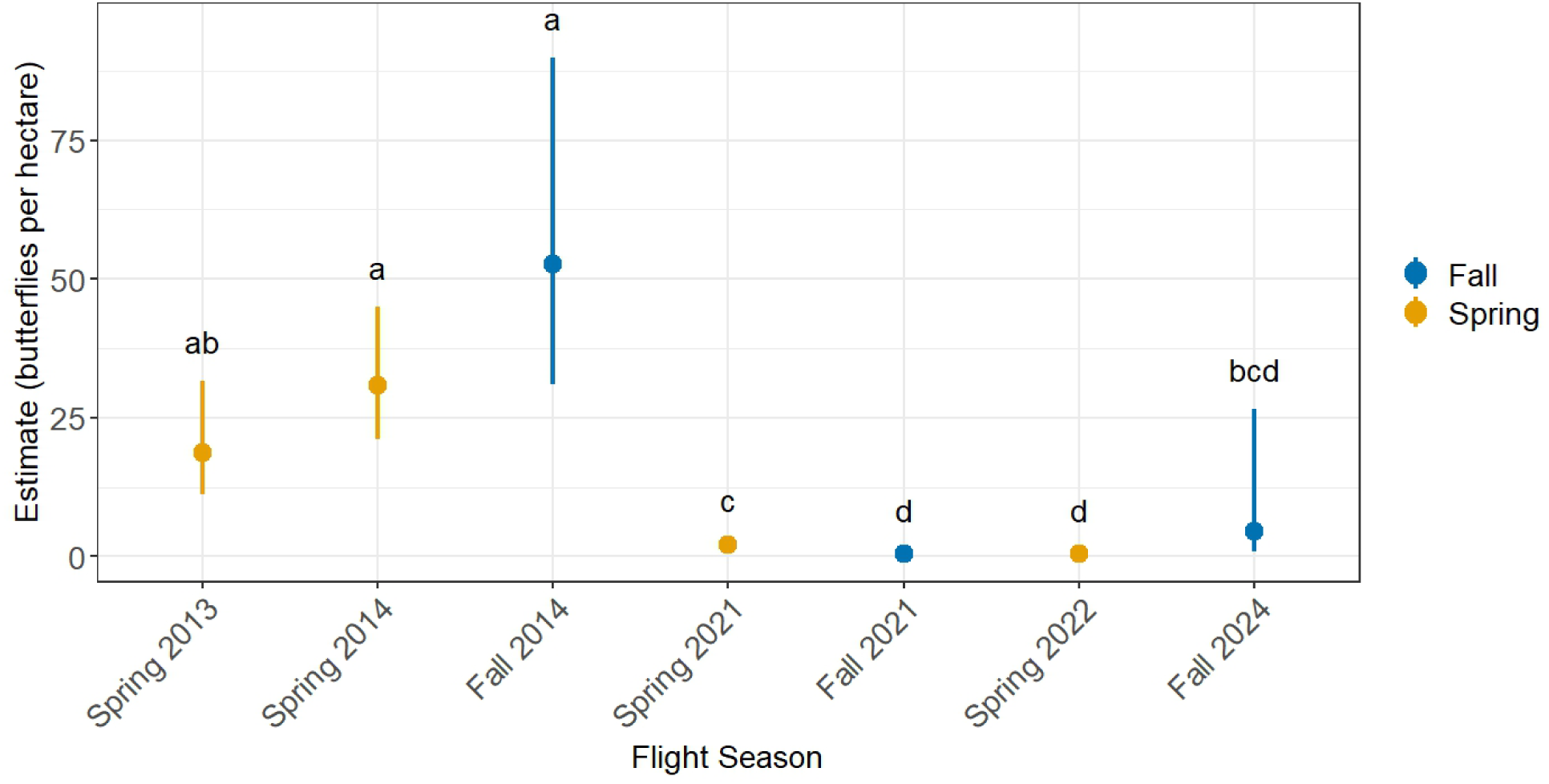
Population estimates for adult butterflies for flight seasons between Spring 2013 and Fall 2024. The points represent the estimates and the whiskers show the 95% confidence intervals. Estimates that share the same letter are not significantly different from one another.

For the period of continuous data collection between February 2021 and February 2022, we observed adult butterflies during every month of year, with peak abundance in February-March 2021 (Figure 6).

**Figure 6:**
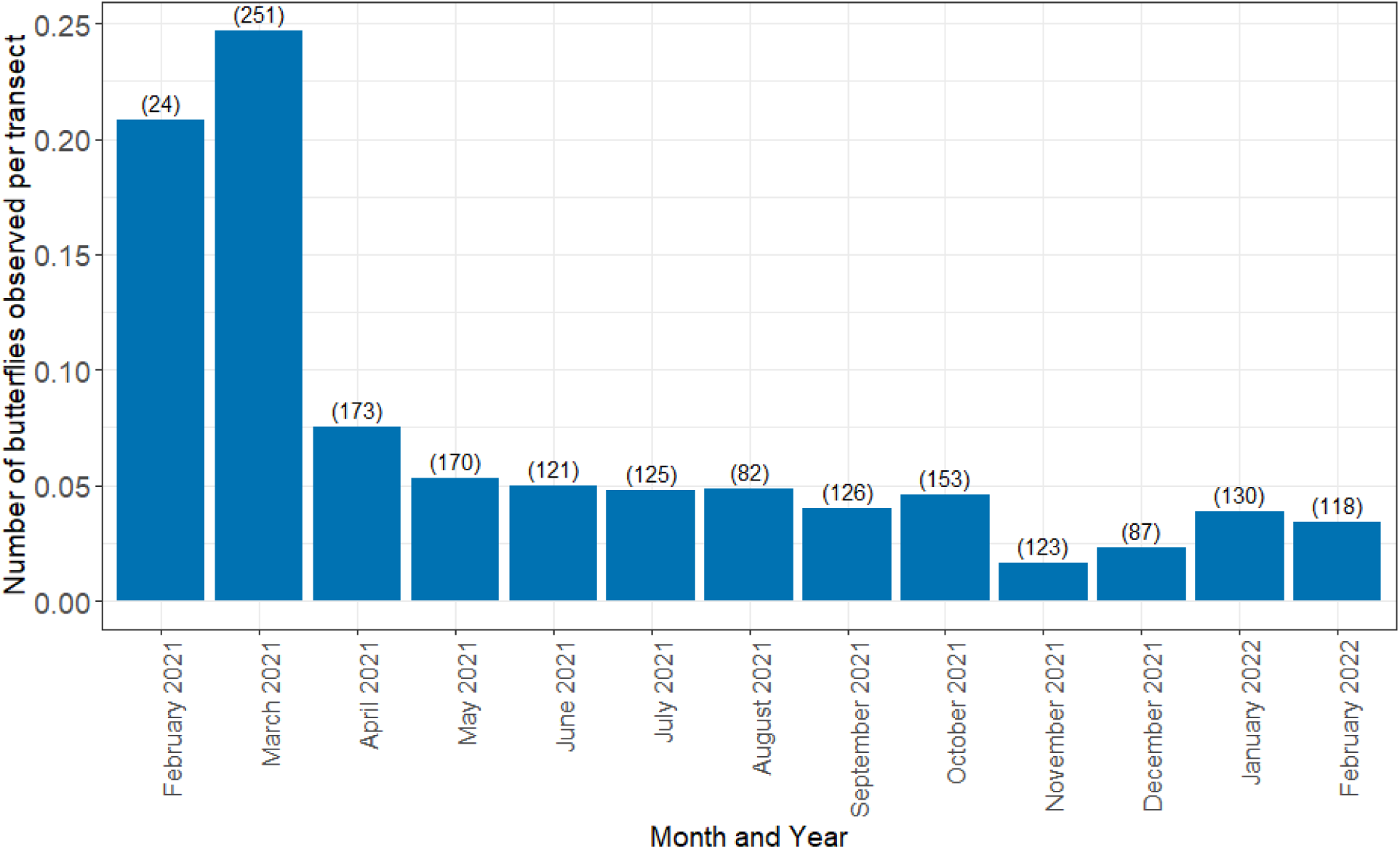
Bar graph showing the number of butterflies observed per transect per month between February 2021 and February 2022. The numbers in parentheses at the top of each bar represent the total number of transects surveyed in each month.

## Discussion

We aimed to examine trends in abundance of both Klot’s sawgrass skipper and its host plant. We found shifts in patterns of host plant abundance between 2013 and 2021 as well as a decrease in the abundance of Klot’s sawgrass skipper between 2013-14 and 2021-24. Our results demonstrate that there is a significant positive relationship between elevation and sawgrass abundance in 2021 where one did not exist in 2013, suggesting that increased salinization is already causing vegetation shifts in this low elevation freshwater ecosystem. The current trend in SLR as measured in Key West, FL is 2.61 mm/yr (± 0.15 mm/yr), for a total increase of 0.27m between 1913 and 2024 and approximately 4.2cm between 2013 and 2021 [43]. Although sawgrass can tolerate short-term pulses in salinity due to tropical cyclones and seasonal hydroperiods, its growth is significantly reduced at salinity levels between 10-20 ppt [44], and germination requires salinity below 5 ppt [45]. Seawater salinity in this region is approximately 35 ppt [46].

Our study is the first to demonstrate changes in the plant community in freshwater wetlands in the Florida Keys, though this phenomenon is well-understood in sawgrass wetlands in the Everglades in mainland South Florida. There, sawgrass communities have shifted geographically due to historical drainage of wetlands, sea level rise, and changing precipitation patterns [47,48]. Shifts in plant communities due to increased salinization is also well-studied in pine rockland forests, a terrestrial ecosystem found adjacent to sawgrass-dominated wetlands. In these forests, a combination of sea level rise and increased salinity during tropical cyclone events led to several measurable impacts including: increased groundwater salinity, an increase in salt-tolerant vegetation [49], and an 80% reduction in pine tree density in these forests on Big Pine Key between 1998 and 2018 [50].

Another source of shifts in vegetation communities in the Lower Florida Keys is a sharp decline in occurrence of fire [51]. Historically, the pine rocklands adjacent to sawgrass wetlands burned approximately every six to nine years [52]. Although fire history has not been well-studied for freshwater sawgrass marshes in the Lower Florida Keys, a natural fire return interval of three to ten years has been established for sawgrass wetlands in the nearby Everglades [53]. Fire suppression in sawgrass wetlands causes increased encroachment by woody shrubs and trees, [54,55] particularly the salt-tolerant buttonwood (Conocarpus erectus) in the Florida Keys [56]. This study did not find an increase in shrub abundance (Figure 3, right panel) despite anecdotal observations of increased shrub cover in sawgrass wetlands. This is likely because the transects with the highest shrub abundance were eliminated from the survey because the high shrub cover makes them nearly impenetrable and difficult to survey on foot. The analysis was done only on transects which were surveyed both in 2013 and 2021, therefore eliminating areas with high levels of shrub encroachment and limiting the scope of our analysis.

Although our butterfly population estimates for Klot’s sawgrass skipper are intermittent, our data indicate likely population declines. Population estimates are consistently much lower in the period from 2021-24 when compared to estimates from 2013-14. It is important to note that high inter-year variability in population size is common in herbivorous insects, often due to fluctuations in weather [57,58] and host and nectar plant quantity and quality [59]. Our population estimates are sensitive to errors, for example issues in estimating distances in the field and variability in observer skill, which can result in large confidence intervals due to limited data [60]. In order to strengthen inferences about the trajectory of Klot’s sawgrass skipper populations, it would be useful to conduct additional population surveys during periods of high observed skipper numbers.

Klot’s sawgrass skipper populations may also be influenced by mosquito control activities on nearby residential properties. Freshwater wetlands in the Lower Florida Keys are contaminated by permethrin due to drift from mosquito control activities on nearby residential properties [61]. Although the dietary risk assessment conducted by Bargar and Hladik concluded that mortality risk due to permethrin contamination of sawgrass was low, their study did not consider sublethal effects of permethrin exposure such as reduced larval growth or decreased adult size [61]. Additionally, there are a number of other pesticides currently used as mosquito adulticides in Klot’s sawgrass skipper habitat [62], including naled [63,64] and malathion [65] which are toxic to butterflies and other non-target invertebrates.

Despite the apparent decreased density of Klot’s sawgrass skipper, its extant range has not significantly contracted; in 2021-22 the butterfly was found on the same islands across the same range as in 2013-14 surveys. Even at very low densities, Klot’s sawgrass skipper has continued to persist in very small patches of habitat. We have observed multiple life stages in isolated roadside patches of habitat no more than a few square meters in size. Based on our survey data, we establish that Klot’s sawgrass skipper is multivoltine and is present year-round as adults in freshwater sawgrass-dominated wetlands (Figure 6), with periods of high abundance in the Spring (Feb-Apr) and in the Fall (October), though some years seem to be missing the Fall flight. Previously, this butterfly was thought to have two generations per year [23]. This presumption was based on the mainland subspecies of this butterfly, likely in the temperate part of its range [38]. This tendency of butterflies and other insects to have more generations per year (increased voltinism) at lower latitudes is common [66,67]. Although the consequences of climate change on butterflies and moths are diverse [68], warming from climate change is leading to increased multivoltinism [69] Sea level rise will increasingly impact the Florida Keys in the near future. Current projections of likely SLR in the Florida Keys are 0.26-0.29m by 2040 and 0.45-0.59m by 2060 [70]. Increased salinity due to SLR is likely to be exacerbated by a projected increased severity of extreme tropical cyclones [71–73]. This region experiences frequent disturbances from tropical cyclones which lead to short-term increases in salinity; the interaction of these disturbances and SLR are likely to further increase salinity [35,74]. Climate change is additionally projected to cause shifts in precipitation, which could lead to decreased precipitation in the wet season but an increased occurrence of extreme precipitation events [75]. Since all of the freshwater in this system is from precipitation [32], precipitation shifts could lead to prolonged periods of high salinity. SLR is already having major impacts in low-lying islands globally [76] including the recent extirpation of an endemic cactus in the Florida Keys due to SLR [77](. Klot’s sawgrass skipper has a high likelihood of joining the ranks of organisms driven to extinction due to SLR – nearly all of its habitat will be rendered unsuitable within several decades with no ability for the habitat to migrate upslope.

Despite the present and future conservation challenges due to climate change, it is vital to manage habitat for as long as possible. In the Florida Keys, this means restoring fire disturbance which increases species’ ability to survive disturbance from hurricanes [74].

Without habitat management actions such as prescribed fire, Klot’s sawgrass skipper risks being adding to the growing list of species that have become extinct or locally extirpated in the Lower Florida Keys for reasons largely unrelated to climate change, as was the was the case with Bartram’s scrub-hairstreak butterfly (Strymon acis bartrami), Florida leafwing butterfly (Anaea troglodyta floridalis), both of which have been apparently extirpated from the Lower Florida Keys due to fire suppression and habitat fragmentation[78,79]. Two other butterflies recently went extinct in South Florida: the Zestos skipper (Epargyreus zestos) and Rockland grass skipper (Hesperia meskei pinocayo) [80,81].

Although the conservation prospects for organisms like Klot’s sawgrass skipper and freshwater marshes in the Florida Keys are bleak in the face of SLR, these most vulnerable taxa and ecological communities present us with a valuable learning opportunity. The problems that these taxa and ecological communities are facing today are a preview of the conservation challenges that will face a broader swath of our globe in the near future. In particular, these canaries in the coalmine present us with living laboratories to understand how adaptive management strategies including ecological restoration can improve resistance and resilience [82].

## Acknowledgements

Many thanks to the field technicians who participated in this project: Erin Binkley, Lauren Breza, Zach Green, Taylor Hunt, Sean Johnson-Bice, Camille Knight, Jessica Padilla, Cody Slaugh, and Adam Emerick. We thank Lydia Soifer, who co-created the plant models presented in this manuscript and the staff of the Florida Keys National Wildlife Refuges Complex including Chris Eggleston, Adam Emerick, Nancy Finley, Heather Stewart, and Kate Watts for supporting this project. Additionally, we thank Dr. Phil Hahn, Dr. Emily Khazan, and Laura Steele for reviewing and improving drafts of this manuscript.

