## Supplementary Material for "Impacts of sea level rise on an endemic butterfly and its freshwater wetland habitat in the Florida Keys"

Supplementary Table 1: Comparison of AICs for various detection functions and series expansions in Distance for the two sampling periods, 2013-14 and 2021-24.

| model | AIC |
| --- | --- |
| data_1314.hnnull | 1021.824 |
| data_1314.hncos | 1016.53 |
| data_1314.hnherm | 1021.824 |
| data_1314.hnpoly | 1021.676 |
| data_1314.hrnull | 1021.302 |
| data_1314.hrcos | 1021.302 |
| data_1314.hrherm | 1009.196 |
| data_1314.hrpoly | 1011.246 |
| data_2124.hnnull | 546.178 |
| data_2124.hncos | 545.651 |
| data_2124.hnherm | 545.935 |
| data_2124.hnpoly | 546.178 |
| data_2124.hrnull | 543.95 |
| data_2124.hrcos | 543.95 |
| data_2124.hrherm | 543.95 |
| data_2124.hrpoly | 543.95 |

Supplementary Table 2: Summary of Distance output for hazard-rate model with Hermite polynomial adjustment for each flight season, with pooled observations across islands.

| Season | Density Estimate | SE | CV | LCL | UCL | DF | GOF Chi-squared | GOF p-value | GOF df |
| --- | --- | --- | --- | --- | --- | --- | --- | --- | --- |
| Spring 2013 | 18.75 | 5.04 | 0.269 | 11.1 | 31.6 | 134.5 | 9.23 | 0.100 | 5 |
| Spring 2014 | 30.86 | 5.94 | 0.192 | 21.2 | 45.0 | 208.8 | 5.28 | 0.382 | 5 |
| Fall 2014 | 52.76 | 14.5 | 0.274 | 30.9 | 90.0 | 101.9 | 32.14 | 5.58E-06 | 5 |
| Spring 2021 | 2.17 | 0.394 | 0.182 | 1.52 | 3.09 | 340.8 | 24.10 | 0.000207 | 5 |
| Fall 2021 | 0.493 | 0.241 | 0.490 | 0.192 | 1.27 | 32.44 | 3.662 | 0.599 | 5 |
| Spring 2022 | 0.382 | 0.263 | 0.689 | 0.108 | 1.35 | 35.69 | NaN | 0.405 | 5 |
| Fall 2024 | 4.55 | 4.58 | 1.00 | 0.780 | 26.6 | 16.8 | 1.889 | 0.864 | 5 |
